# Complex heterochrony underlies the evolution of hermaphrodite self-fertility and sex allocation in experimental *C. elegans* populations

**DOI:** 10.1101/047811

**Authors:** Nausicaa Poullet, Anne Vielle, Clotilde Gimond, Sara Carvalho, Henrique Teotónio, Christian Braendle

## Abstract

Hermaphroditic organisms are common both in plants and animals, and have served as key models to study the evolution of sex allocation. Despite extensive past research, the specific developmental mechanisms by which hermaphrodite sex allocation can evolve remain largely unknown. To address this problem, we here use experimental evolution of *Caenorhabditis elegans* hermaphrodite-male populations to directly quantify changes in germline and somatic development that underlie adaptive shifts in hermaphrodite sex allocation associated with the evolution of improved self-fertility. Specifically, we test whether the evolution of hermaphrodite sex allocation is due to heterochrony, i.e. evolutionary changes in the relative timing of developmental processes.

We show that the experimental evolution of improved hermaphrodite self-fertility occurred through complex modification of a suite of developmental and reproductive traits: increased sperm production, accelerated oogenesis and ovulation rates, and increased embryo retention *in utero*. The experimental evolution of increased sperm production delayed entry into oogenesis – as expected, given the sequentially coupled production of spermatogenesis and oogenesis. Surprisingly, however, delayed oogenesis onset did not delay reproductive maturity, nor did it trade-off with gamete or embryo size. Comparing developmental dynamics of germline and soma indicates that the evolution of increased sperm production did not delay reproductive maturity due to a globally accelerated larval development during the period of spermatogenesis.

We conclude that the integration of multiple heterochronic events in gametogenesis and soma can explain the experimental evolution of hermaphrodite sex allocation and self-fertility. Our results thus support the idea that heterochrony can represent a specific mechanism that explains the maintenance of partial selfing in natural populations with mixed reproduction modes and different forms of hermaphroditism. More generally, our results provide a quantitative perspective on how natural selection can operate on developmental characters – and their integration – during the evolution of life history at the population level.

## 1. Introduction

How organisms allocate their resources to different sexual functions is central to reproductive fitness, so that much theoretical and empirical research in evolutionary biology focuses on sex allocation. In many taxa, sex allocation may be realized through the maintenance of separate sexes, each specialized in female versus male reproductive functions. Alternatively, species may show co-sexuality, i.e. hermaphroditism where individuals perform both female and male reproductive functions. Self-fertilization (selfing) through hermaphrodites is common in animals and plants (Stebbins 1957; Jarne and Auld 2006; Weeks et al. 2006; Barrett 2008), and a central question is thus how natural and sexual selection can act on the allocation to male versus female function within individuals of hermaphroditic organisms.

Fisher’s insight that negative frequency-dependent selection determines the evolution of sex ratios (Fisher 1930) can be used to understand how selection acts on hermaphrodite sex allocation. Specifically, the marginal fitness returns of alleles responsible for conflicting sexual functions in hermaphrodites are expected to be minimized during evolution (Charlesworth and Charlesworth 1981; Charnov 1982; Lloyd 1984). When hermaphrodites function as one sex during early-life and another sex late in life, the direction of sex change during development will depend on resource availability and the timing of resource allocation towards reproduction versus survival (Brunet 1992; Kinkhammer et al. 1997). On the other hand, if hermaphrodites have simultaneous sex functions, availability of mates and breeding seasonality may determine how natural and sexual selection will operate on developmental time to maturity (Schärer 2009; Escobar et al. 2011). When hermaphrodites predominantly self, the even allocation of resources towards spermatogenesis and oogenesis should be favoured (Charnov 1987). As reproduction is usually oocyte-limited for most organisms, female-biased sex allocation is expected to be favoured under predominant selfing (Weeks 2012).

Some of these theoretical predictions concerning the evolution of hermaphrodite sex allocation have been experimentally examined by manipulating mating or cross-pollination opportunities (Tian-Bi et al. 2008; Dorken and Pannell 2009; Roels and Kelly 2011), the level of resources available for reproduction and survival (Mazer et al. 2003; Case and Ashman 2007), and by manipulating the reproductive system of the populations under study (Teotonio et al. 2012; Carvalho et al. 2014b; Theologidis et al. 2014). Despite these studies, however, it remains mostly unknown which developmental changes underlie the evolution of hermaphrodite sex allocation under selfing and/or outcrossing (Li and Johnston 2010; Roels and Kelly 2011; Noel et al. 2016). In general, it is evident that the evolution of hermaphrodite sex allocation may occur through selection on the timing and rates of gametogenesis and other developmental processes associated with sexual functions (Alberch et al. 1979; Lord and Hill 1987; Li and Johnston 2000; Mazer et al. 2004). Such heterochrony – defined here as an evolutionary change in the relative timing or relative rates of developmental processes (Alberch et al. 1979; Li and Johnston 2000) – has been suggested to be a common mechanism underlying the evolution of hermaphrodite outcrossing rates and evolutionary transitions between selfing and outcrossing.

Evolutionary transitions to selfing have been correlated with the evolution of developmental time to maturity (Lord and Hill 1987; Weeks et al. 2006; Tian-Bi et al. 2008; Escobar et al. 2011) and, in the case of angiosperms, with differential timings and rates of within flower germline and somatic tissue development (Darwin 1877; Harder and Barrett 2006; Li and Johnston 2010; Geuten and Coenen 2013). Most of these studies have inferred heterochrony based on comparative analysis, and we therefore lack direct measurements of evolutionary responses in relevant developmental processes and their timing during the evolution of hermaphrodite sex allocation in experimentally defined contexts. Here we use experimental evolution of *Caenorhabditis elegans* to quantify heterochrony in germline and somatic development that may underlie adaptive shifts in hermaphrodite sex allocation associated with the evolution of improved self-fertility.

The nematode *C. elegans* exhibits androdioecy, a reproductive system consisting of self-compatible hermaphrodites and males (Maupas 1900). In natural populations, *C. elegans* hermaphrodites primarily self and males are rarely found (Barriere and Felix 2005). Unlike most hermaphroditic animals and plants, *C. elegans* hermaphrodites cannot mate with each other and thus exclusively function as females when outcrossing occurs through male mating. The “male” function of *C. elegans* hermaphrodites is thus limited to the process of spermatogenesis and represents an exclusive component of selfing, being better termed “hermaphroditic” or “selfing” function. In contrast, female functions of *C. elegans* hermaphrodites, encompassing oogenesis, ovulation, mating and egg-laying, are common to both selfing and outcrossing. The inability of hermaphrodites to fertilize each other renders *C. elegans* particularly useful as selfing versus outcrossing rates can be unambiguously determined (Stewart and Phillips 2002).

*C. elegans* hermaphrodites are protandrous, with early germ cells differentiating into sperm, followed by an irreversible switch to oogenesis around the larval-adult moult transition (Hirsh et al. 1976). Due to this sequential nature of gametogenesis, allowing potentially continued oogenesis until death, selfing hermaphrodites are self-sperm limited under most demographic circumstances (Barker 1992; Cutter 2004). Importantly, due to the sequential coupling of spermatogenesis and oogenesis, increasing sperm production (by means of a prolonged period of spermatogenesis) causes a delay in the onset of oogenesis and reproductive maturity, thus generating a fitness trade-off between generation time and lifetime fecundity (Hodgkin and Barnes 1991; Cutter 2004). The timing of the switch from spermatogenesis to oogenesis represents a direct regulatory mechanism of hermaphrodite sex allocation, so that timing shifts of this switch represents an obvious candidate mechanism underlying the evolution of sex allocation in *C. elegans* hermaphrodites.

While outcrossing is rare in natural *C. elegans* populations, intermediate frequencies of outcrossing can be maintained when populations with standing genetic variation for male outcrossing ability evolve in experimental environments (Anderson et al. 2010; Teotonio et al. 2012; Carvalho et al. 2014b). In the experiments of Teotonio et al. (2012) partial selfing was stably maintained at 50% over 100 generations of adaptation to a laboratory environment characterized by discrete and non-overlapping 96-hour life cycles at constant and high population sizes (N=10^4^). The maintenance of partial selfing in these populations can be explained by a counterbalance of evolutionary improvements of male outcrossing ability versus hermaphrodite self-fertility (Carvalho et al. 2014a; Carvalho et al. 2014b). The evolution of improved hermaphrodite self-fertility, i.e. an increase of reproduction at earlier ages relative to total reproduction, is consistent with adaptation to the imposed shortened life cycle during experimental evolution, but may further reflect an evolutionary response that promotes selfing and/or limits outcrossing (LaMunyon and Ward 1998; Carvalho et al. 2014b). Interestingly, evolved hermaphrodites did not show increased relative reproduction at an early age when fertilized by males (LaMunyon and Ward 1998; Carvalho et al. 2014b). Since the female function in *C. elegans* hermaphrodites is common to both selfing and outcrossing, this result suggests that hermaphrodite spermatogenesis and sperm production - as the exclusive selfing developmental component - must have been improved during evolution (LaMunyon and Ward 1998; Carvalho et al. 2014b). Consequently, because increased sperm production delays reproductive maturity of selfing *C. elegans* hermaphrodites (Hodgkin and Barnes 1991; Cutter 2004), the observation that evolved hermaphrodites also showed increased early-life fertility appears paradoxical: how can evolution of increased hermaphrodite sperm production be developmentally accommodated without causing a delay in reproduction, i.e. decreased early-life fertility? Diverse evolutionary changes in reproductive development, including heterochrony, could explain this seemingly contradictory finding; however, the specific changes in hermaphrodite sex allocation and developmental timing that occurred during these evolution experiments remain to be determined.

The above experimental observations motivated the present study to directly measure changes in *C. elegans* hermaphrodite development that underlie the evolution of improved self-fertility and apparently associated changes in hermaphrodite sex allocation. Using the same experimental populations (Carvalho et al. 2014a; Carvalho et al. 2014b) (Fig. 1A), we quantified in parallel the relative differences in developmental and reproductive traits between selfing hermaphrodites of ancestral (generation 0) versus evolved populations (generation 100), in the context of the 96-hour life cycle imposed during experimental evolution. We followed the dynamics of hermaphrodite germline and somatic development throughout the life cycle to quantify somatic developmental timing, germline progression and differentiation, transition from spermatogenesis and oogenesis, ovulation and gamete size (Fig. 1B,C). In parallel, these developmental phenotypes were integrated with measurements of relevant reproductive output traits, including the timing of reproductive maturity, offspring number and size, and reproductive schedules. Using these measures, we ask whether heterochrony in germline and/or soma was responsible for the evolution of improved hermaphrodite self-fertility, and as a consequence, whether such heterochrony can explain the observed adaptive maintenance of partial selfing in experimental populations.

**Figure 1.**
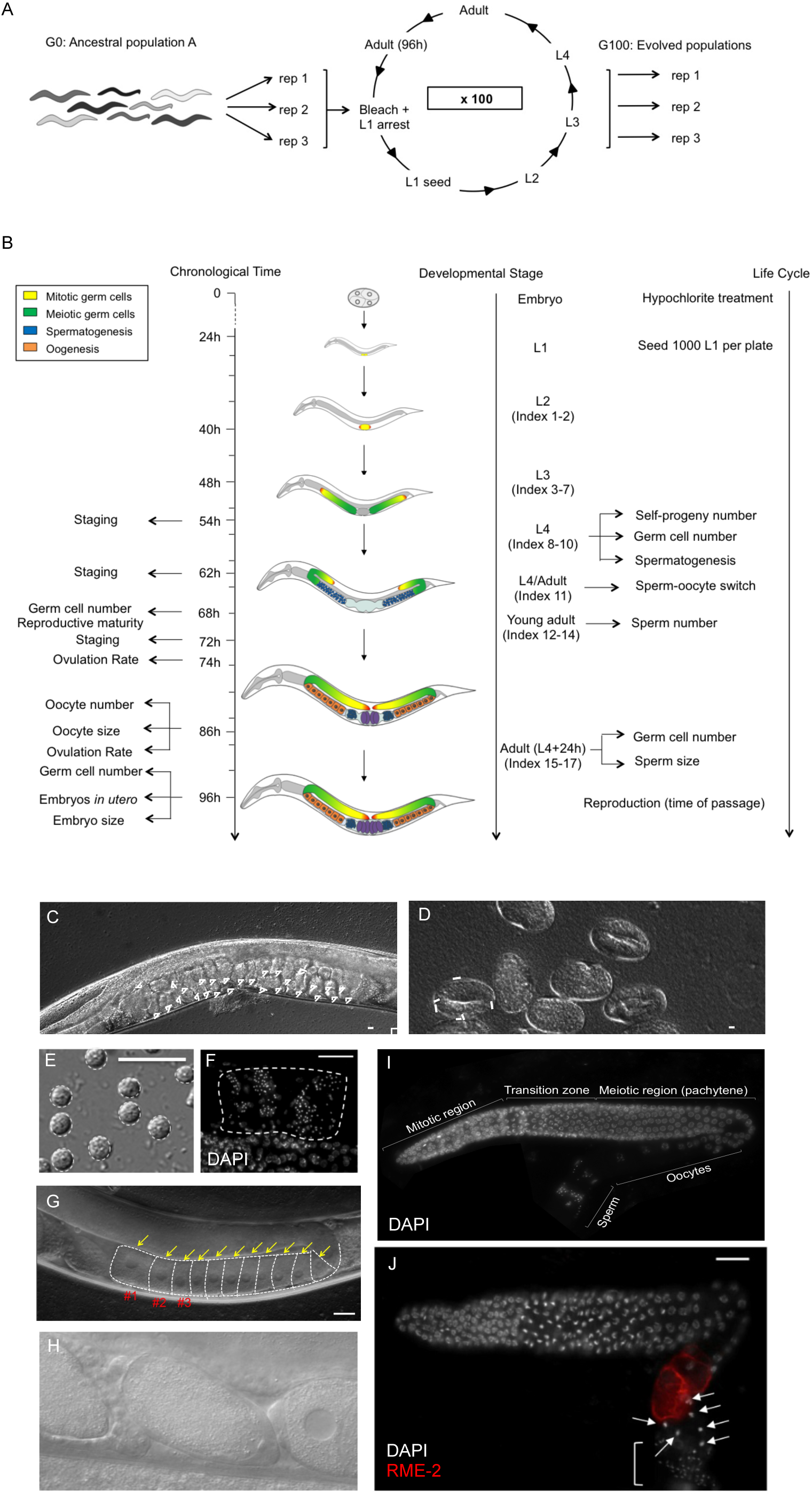
Life-cycle during experimental evolution, hermaphrodite germline development, and experimental design. **(A)** Outline of the experimental evolution assay and 96-hour life cycle (Teotonio et al. 2012). The ancestral androdioecious population G0 was derived from a funnel pairwise cross of 16 *C. elegans* wild isolates. Starting from this population, several independent replicate populations were cultured for 100 generations at 20°C following the same 96-hour life cycle (Fig. 1A) (Teotonio et al. 2012), and three of these replicate populations (G100, rep 1-3) were examined in this study. The time of passage between generations is defined as the life cycle end time point of adults at 96h, which are harvested to collect embryos, defining the starting time point of the life cycle (0h) of the subsequent generation. Collected embryos were maintained in liquid medium without food for 24h during which they hatch into L1 larvae that arrest development. At this time point (24h), age-synchronized L1 cohorts were seeded onto agar plates with bacterial food at a constant population size (N=1000/plate). Populations were then allowed to develop and reproduce for 72h until the time of passage (96h) at which time individuals were washed off plates, treated with a bleach-hypochlorite solution to harvest embryos, initiating the next generation. **(B)** Timeline of phenotyping assays performed in this study. All phenotypes were measured in the same life cycle as imposed during the 100 generations of experimental evolution (Teotonio et al. 2012). Phenotypes were concurrently scored in the ancestral population and three evolved replication populations. All phenotypes – except ovulation rates – were measured in parallel in a single experiment (see Supplementary materials). Different phenotypes were quantified in chronologically staged hermaphrodites at different time points (left) and in hermaphrodites of morphologically defined, distinct developmental stages (right). The drawings illustrate the developmental progression of the hermaphrodite through the four larval (L1 to L4) and adult stages during the life cycle. Timing of developmental events is based on observations of the ancestral population. Different colours depict mitosis-meiosis progression and sperm-oocyte allocation in the developing hermaphrodite germline, based on published information (Kimble 1988; Kimble and Crittenden 2007). Distal Tip Cells are indicated in red. **(C-J)** Illustration of a subset of scored phenotypes (based on individuals from the ancestral population). **(C)** DIC image of an adult hermaphrodite at 96h of the life cycle. Embryos in utero are outlined in white. Scale bar: 20p.m. **(D)** DIC image of embryos collected at 96h of the life cycle to estimate embryo size. Scale bar: 20μm **(E)** DIC image of dissected spermatids of an early adult hermaphrodite, used for estimating sperm size. Scale bar: 10μm. **(F)** Microscopy image of a DAPI-stained hermaphrodite at the early adult stage, outlining spermatids in spermatheca and uterus. Scale bar: 20μm. **(G)** DIC image of the posterior gonad arm in an adult hermaphrodite at 86h of the life cycle. Enlarged oocytes developing after the germline loop region are in indicated with yellow arrows. The three most proximal oocytes, labelled 1-3, were used to estimate oocyte size. Scale bar: 20μm **(H)** Nomarski images of spermatheca (with uterus on left, proximal germline on right) in an ovulating hermaphrodite (74h of life cycle). **(I)** Microscopy image of an extruded DAPI-stained gonad arm of an adult hermaphrodite (distal-proximal axis oriented from left to right). Mitotic and meiotic (pachytene) nuclei can be distinguished using nuclear morphology. The transition zone delineates the entry into meiosis, apparent by crescent-shaped nuclei (Francis et al. 1995; Crittenden and Kimble 2008). Scale bar: 20μm **(J)** Microscopy image of dissected gonad arm of a hermaphrodite at the L4/adult stage (ancestral population), stained with DAPI (white) and RME-2 antibody (red). This protocol was used to infer measures on entry into spermatogenesis and oogenesis, respectively. Mature spermatids are located in the most proximal zone of the gonad, delineated by the white bracket; distinct, large nuclei of primary spermatocytes are indicated by arrows. Staining with RME-2 indicates presence of differentiating oocytes in the proximal germline. Scale bar: 20μm

## 2. Materials and Methods

### (a) Experimental evolution life-cycle and experimental design

All results reported here are based on experimental evolution of androdioecious *C. elegans* populations in the context of a 96-hour experimental life cycle (Fig. 1A). The ancestral population (G0) was constructed by the successive pairwise inter-crossing of 16 isogenic wild isolates. Starting from this population, several independent replicate populations were cultured for 100 generations at 20°C and 80% RH under a discrete time, 96-hour non-overlapping life cycle with a population census size of 10^4^ individuals (from first larval stage to adulthood), and effective population sizes of about 10^3^ (Teotonio et al. 2012; Chelo and Teotonio 2013). Three of these G100 replicate populations (labelled A4-6) were examined in the present study. After 100 generations, populations continued to be genetically diverse, with average genome-wide heterozygosity of about 0.3 (Chelo and Teotonio 2013). At 24h of the life-cycle, 10^4^ L1-staged worms were seeded in ten Petri plates with NGM-lite agar (US Biological) and covered with an *E. coli* HT115 lawn that served as *ad libitum* food until reproduction. 72h±2h later, corresponding to 96h/0h (end/starting point) of the life cycle, adults from all plates were mixed and killed using a hypochlorite “bleach” solution and embryos harvested in a M9 solution (Stiernagle 1999). After 24h±2h, hatched embryos became starvation-arrested L1s, which upon appropriate density estimation are seeded into fresh plates to constitute the following generation (Fig. 1A). Most evolutionary trait changes are expected to be due to the sorting of standing genetic variation (Teotonio et al. 2009; Denver et al. 2010). Further information on the experimental evolution design is presented in Fig. 1A and in Teotonio et al. (2012).

### (b) Life history and developmental assays

For all the assays reported here, the ancestral (G0) and the three evolved (G100) populations were thawed from frozen −80°C samples (each with N>10^3^ individuals) and passaged in parallel for two generations under the standard conditions of experimental evolution. This allowed us to concurrently assay ancestral and evolved populations, control for transgenerational environmental effects and account for potential lab biases.

To confirm the adaptive significance of partial selfing, we determined reproductive schedules of individual selfed hermaphrodites as in Carvalho et al. (2014b). 56h L4 larvae were sampled and transferred to fresh plates every 24h until offspring production ceased. The number of hatched larvae (L1-L3 stages) was scored, for an average of 82.5±5SD individuals per population sample. At 96h of the life cycle, embryo number *in utero* was quantified in randomly picked hermaphrodites (Fig. 1C), for 100 observations per population. Embryo size was quantified at 96h for 200 embryos per population.

Hermaphrodite spermatid area was scored as a proxy for sperm size of dissected young adults (LaMunyon and Ward 1999) that had been isolated 24 hours before the L4 stage to prevent mating with males. An average of 12.5±8.0SD sperm in each of 28-40 individuals were measured per population. Sperm number in young adult hermaphrodites that had been isolated at the L4 stage were measured for an average of 76.3±3.3SD individuals per population.

To characterize spermatogenesis and the timing of the sperm-oocyte switch, at the mid-L4 stage, we examined the proportion of individuals entering spermatogenesis, as inferred from the presence of primary/secondary spermatocytes and mature sperm (an average of 62.5±4.2SD individuals per population was measured). This was done using whole-body DAPI stains. At the L4/adult transition (moult), we scored each individual according to the index: 1: presence of mature spermatids; 2: presence of spermatids and expression of an early oogenesis marker, RME-2 (Grant and Hirsh 1999); and 3: presence of spermatids, RME-2 expression and presence of enlarged diakinesis-stage oocytes. An average of 41±3.5SD individuals per population were measured. Because of the antibody staining, these observations were made using dissected DAPI-stained gonad arms.

We estimated ovulation rates in randomly selected hermaphrodites by scoring the presence or absence of oocytes in the spermatheca (Fig. 1H). 255 individuals (at 74h) and 355 individuals (at 86h) were examined per population. At 86h, for an average of 45.8±0.9SD individuals per population, we counted the number of enlarged oocytes at diplotene and diakinesis of Prophase I, developing in the loop region (McCarter et al. 1999) (Fig. 1G), and we measured the area of the three most proximal oocytes adjacent to the spermatheca (McCarter et al. 1999) (Fig. 1G). We quantified the number of germ cell precursors in hermaphrodites in two somatic stages, mid-L4 and L4 to adult moult, and two chronological stages: 68h and 96h. For all individuals, a single gonad arm was extruded and DAPI-stained to count total germ cell number and to distinguish mitotic versus meiotic germ cell fate, following previously described protocols (Poullet et al. 2015) (Fig. 1I). For each of the four examined developmental stages, analysis included on average 43.4±9.0SD individuals per population.

At 54h, 62h and 72h of the life cycle, an average of 192±24SD randomly selected hermaphrodites per population was scored for their developmental stage, according to supplementary Table S1. Stages of index 1-11 were based on somatic markers and stages of index 12-17 on reproductive markers. For analysis, we first calculated the frequency-weighted average of the numerical index.

Detailed assay descriptions can be found in supplementary materials and methods and Fig.1.

### (c) Statistical analysis

Data was analysed with a linear mixed effects models (LMM) or generalized linear mixed effect models (GLMM) and REML estimation methods (Pinheiro and Bates 2000), to test for significant evolutionary responses, by taking generation as a fixed two level factor and individual (if needed be) nested within population nested within block as random factors. Taking population as a random factor irrespective of generation accounts for the effects of genetic drift and other historical accidents during evolution, while being conservative as the ancestral population variation is considered. Progeny number until 96h over lifetime progeny, and counts of individuals that had entered in spermatogenesis versus those that had not, were analysed with GLMM assuming binomial errors. Embryo number *in utero* and number of oocytes were analysed with GLMM assuming Poisson errors. All other traits were modelled with LMM. Model residuals were tested for normality with Shapiro-Wilk tests, when evolutionary responses were significant. In the cases of spermatogenesis entry, sperm-oocyte transition and reproductive maturity in staged hermaphrodites, outlier removal (residuals >2.5) and data transformation were unsuccessful in normalizing residuals. In these cases, LMM or GLMM models were performed without considering generation as a factor, and likelihood ratio tests (LRtest) done to test whether inclusion of generation improved the models’ fit. LRtest significance was determined assuming a Chi-squared distribution with one degree of freedom, and also by parametric bootstrapping (PBtest) (Halekoh and Højsgaard 2014). For all other traits, planned contrasts among generations were done with Student t tests with the effective number of degrees of freedom being calculated with the Kenward-Roger approximation (Halekoh and Højsgaard 2014; Lenth 2014). Even if evolutionary responses were not significant, we present in all plots the least-square estimates from the models. The packages *stats, lme4, lsmeans* and *pkbrtest* in R were used for computation (R Development Core Team 2013).

## 3. Results

### (a) Evolution of hermaphrodite self-fertility and reproductive schedule

We first compared reproductive schedules of selfing hermaphrodites between ancestral and evolved hermaphrodites to confirm the adaptive significance of partial selfing during experimental evolution (Carvalho et al. 2014b). Consistent with adaptation to the experimental 96-hour life cycle, evolved hermaphrodites produced a higher number of selfed offspring until 96h (time of passage) when concurrently compared to ancestral hermaphrodites (Fig. 2A; one-tailed t_5.4_ P=0.03), while lifetime number of offspring was unchanged (Fig. 2B; t_5.1_ P=0.67). There was thus a shift towards increased early self-reproduction with experimental evolution as expected with adaptation (Williams 1957; Anderson et al. 2011). We further found that, at 96h, evolved hermaphrodites exhibited a much higher number of embryos *in utero* (Fig. 2C; t_4.2_ P<0.01), indicative of strong embryo retention. In contrast, embryo size did not change during evolution (Fig. 2D; t_5_ P=0.4). Together, these findings suggest that the evolution of improved hermaphrodite reproduction was partly due to the evolution of increased embryo retention, probably as a direct response to the hypochlorite treatment imposed at 96h of the life cycle, throughout experimental evolution and to which only embryos survive (Stiernagle 1999; Teotonio et al. 2012).

**Figure 2.**
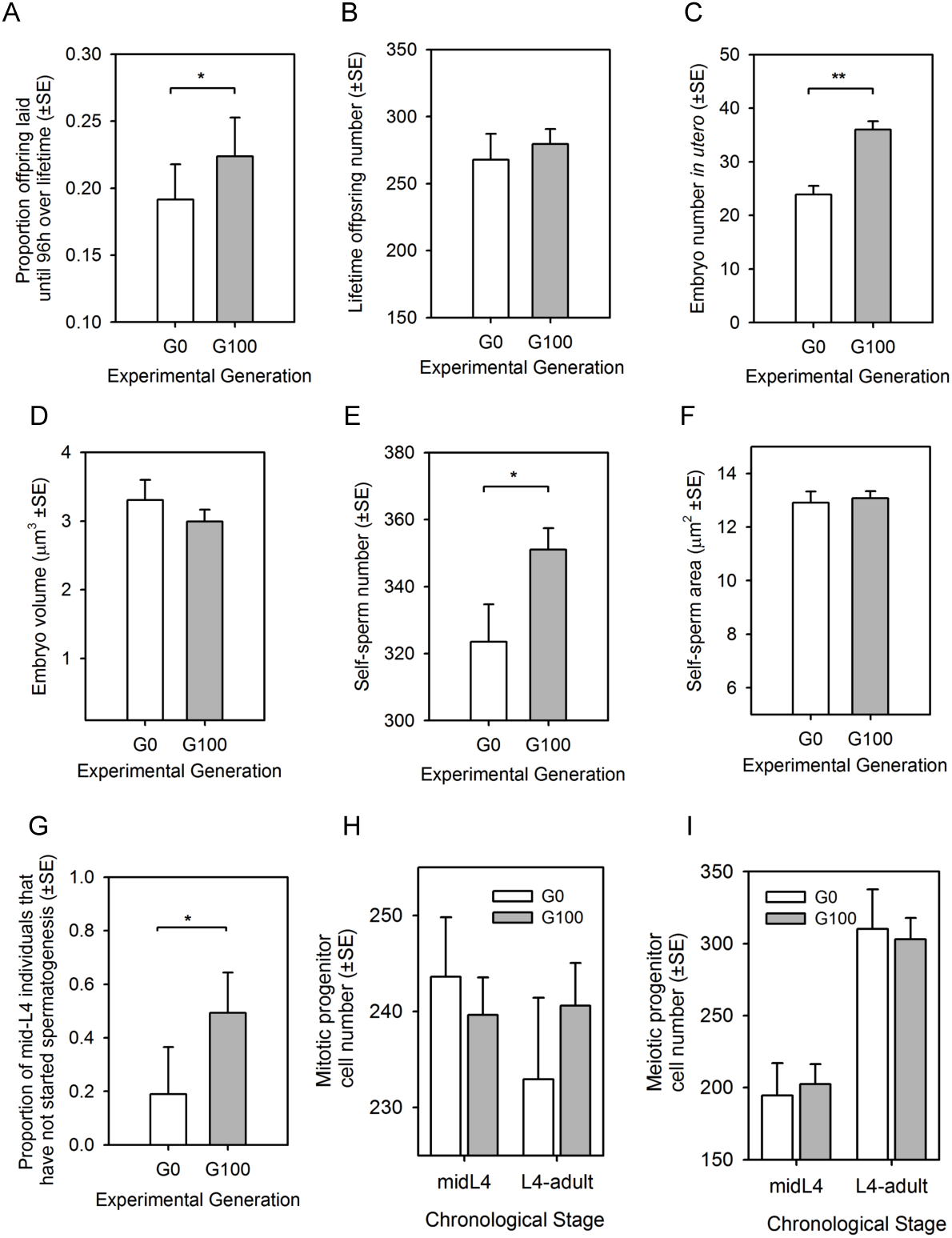
Evolution of reproduction, embryonic traits, sperm traits and germ cell progenitors. **(A)** Reproductive schedules of self-fertilization hermaphrodites in ancestral and evolved populations. Evolved population showed a significant shift towards earlier reproduction. **(B)** Lack of experimental evolution responses in lifetime offspring production of selfed hermaphrodites. **(C)** Number of embryos in utero at 96h, the time of passage during experimental evolution. There was an increase of embryo number in utero during experimental evolution. **(D)** Embryo size (volume, μm3 x 10-4) measured at 96h did not differ between ancestral and evolved populations. **(E)** Hermaphrodite self-sperm number significantly increased with experimental evolution. **(F)** Hermaphrodite self-sperm size (cross-sectional area, μm2) did not significantly differ between evolved and ancestral populations. **(G)** Evolved populations showed a significantly lower proportion of mid-L4 individuals that had initiated sperm differentiation (formation of primary spermatocytes). **(H)** Mitotic germ cell numbers (per gonad arm) for somatically staged individuals (mid-L4 and L4/adult moult). **(I)** Meiotic (pachytene) germ cell numbers (per gonad arm) for somatically staged individuals (mid-L4 and L4/adult moult). Linear mixed effects and generalized linear mixed effects models used to test for differences among generation 0 (G0) and generation 100 (G100): * P-values<0.05, ** for P-values<0.01, *** for P-values<0.001. Means and error least square estimates are shown. See main text for summary statistics.

### (b) Evolution of spermatogenesis

Hermaphrodites of evolved populations produced significantly more sperm than hermaphrodites of ancestral populations (Fig. 2E; one-tailed t_5_ P=0.04); in contrast, hermaphrodite sperm size did not evolve (Fig. 2F; t_4.3_ P=0.75). In somatically-staged mid-L4 larvae, we further found that the germline of all individuals in the ancestral population contained primary spermatocytes and/or mature spermatids whilst a significant proportion of individuals in evolved populations had not yet started spermatogenesis (Fig. 2G; one-tailed t_4.1_ P=0.03, LRtest P=0.02, PBtest P=0.05). Evolution of increased self-sperm production was thus accompanied by a delayed entry into spermatogenesis relative to somatic developmental stage. This result suggests that evolution of increased sperm production caused a delayed entry into spermatogenesis, so that the increased allocation of precursors to the sperm fate delays the onset of spermatocyte formation. However, the number of germ cell progenitors in mid-L4 larvae, when spermatogenesis is just beginning did not differ between evolved and ancestral hermaphrodites (Fig. 2H,I;, mitotic: t_61_ P=0.57, meiotic: t_15.3_ P=0.77).

Similarly, at the L4-adult moult, the numbers of precursor cells in the mitotic or in the meiotic phases did not differ between evolved and ancestral hermaphrodites (Fig. 2H,I; mitotic: t_144_ P=0.6, meiotic: t_30_ P=0.82).

### (c) Evolution of the sperm-oocyte switch

Without the need to invoke changes in the rates of spermatogenesis, the evolution of increased hermaphrodite sperm production (and delayed entry into spermatogenesis) could have resulted in a delay of the transition between spermatogenesis and oogenesis as they are sequentially connected (Kimble 1988; Kimble and Crittenden 2007). To confirm this scenario, we inferred the timing of the sperm-oocyte switch using an index that takes into account three distinct events of gamete differentiation and maturation in somatically-staged hermaphrodites at the L4/adult moult: presence of mature spermatids, expression of the early-oogenesis marker RME-2 (Grant and Hirsh 1999), and presence of oocytes (diplotene-diakinesis stages) in the proximal gonad arm. As expected, we found a decrease in the index and thus what can be interpreted as a small but significant delay in the sperm-oocyte switch and delayed entry into oogenesis, relative to soma (Fig. 3A; one-tailed t_4.1_ P=0.03, LRtest P=0.02, PBtest P=0.05). This result is thus consistent with the observations of delayed entry into spermatogenesis and prolonged spermatogenesis in evolved hermaphrodites.

**Figure 3.**
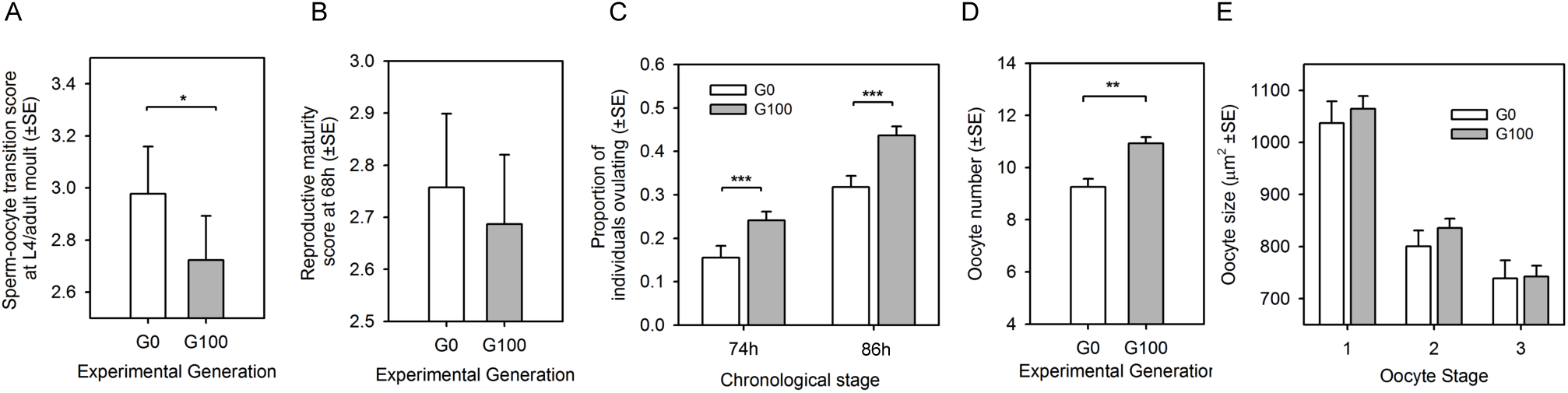
Evolution of the sperm-oocyte switch, oogenesis, ovulation, and germline progression during adulthood. **(A)** Evolved populations showed a significant delay of the sperm-oocyte decision relative to somatic developmental stage (L4/adult moult), based on an index scoring presence of mature spermatids, expression of the early-oogenesis marker RME-2, and presence of oocytes (diplotene-diakinesis stages) in the proximal gonad arm. **(B)** At 68h of the life cycle, ancestral and evolved populations did not differ in reproductive maturity (presence of oocytes and/or embryos in uterus). **(C)** The number of oocytes was significantly increased at 74h after experimental evolution. **(D)** Ovulation rates at 74h and 86h of the life cycle were significantly increased after experimental evolution. **(E)** Oocyte size (area, μm2 x 10-4) did not differ between ancestral and evolved populations. Linear mixed effects and generalized linear mixed effects models used to test for differences among generation 0 (G0) and generation 100 (G100): * P-values<0.05, ** for P-values<0.01, *** for P-values<0.001. Means and error least square estimates are shown. See main text for summary statistics.

### (d) Evolution of oogenesis and ovulation

Contrary to expectations (Hodgkin and Barnes 1991; Murray and Cutter 2011), the observed delay in the switch between spermatogenesis and oogenesis did not result in a delay in reaching reproductive maturity since the proportion of individuals with oocytes and first embryos was comparable between evolved and ancestral populations at 68h of the life cycle (Fig. 3B; one-tailed t_5.6_ P=0.17, LRtest P=0.29, Pbtest P=0.27).

Measures of ovulation rates in chronologically-staged hermaphrodites at 74h and 86h indicated a significant acceleration of ovulation in evolved hermaphrodites (Fig. 3C; 74h: t_24.2_ P<0.01; 86h: t_21_ P<0.001). At 86h, evolved hermaphrodites also possessed higher numbers of oocytes relative to ancestral hermaphrodites (Fig. 3D; t_4.8_ P<0.01), while oocyte size did not evolve (Fig. 3E; t_4.9_ P=0.5). Thus, despite accelerated ovulation, more oocytes were present in gonads of evolved hermaphrodites, which indicates that oogenesis became significantly faster with evolution. Accelerated oogenesis and ovulation, despite a concurrently increased sperm production and developmental delay in the sperm-oocyte switch may therefore have contributed to the evolution of increased self-reproduction at earlier ages.

As for earlier developmental stages, the number of mitotic or meiotic germ cell precursors was unaffected with evolution in chronologically staged hermaphrodites at 68h (Fig. S2; mitotic: t_79.9_ P=0.57 meiotic: t_17.9_ P=0.24;), and at 96h, the time of reproduction (Fig. S2, mitotic: t_57.4_ P=0.55, meiotic: t_15_ P=0.18). Therefore, although evolved populations displayed faster oogenesis and ovulation, we did not detect a correlated response in oogenic precursor numbers, e.g. a reduction of meiotic precursors in evolved populations. This discrepancy may reflect accelerated germ cell proliferation and their meiotic differentiation in evolved populations, which could compensate for the observed increases in oocyte numbers.

### (e) Evolution of developmental timing in soma and germline

To understand how germline progression and differentiation integrates with developmental timing across the fixed 96h life-cycle, we quantified somatic progression and reproductive maturation by chronological staging of individuals at 54h, 62h and 72h of the life cycle (Fig. 1B, Table S1). Larval stages at 54h and 62h were inferred from analysis of vulval cell lineages and morphogenesis (Sulston and Horvitz 1977; Sternberg and Horvitz 1986). Results show that there was no evolutionary response at 54h but evolved populations were developmentally advanced at the intermediate 62h time point relative to the ancestral population (Fig. 4A; 54h: t_16.3_ P=0.92; 62h: t_16.3_ P<0.01). Specifically, at 54h, evolved and ancestral hermaphrodites were mostly late L3 larvae, but at 62h, most hermaphrodites of evolved populations were at the L4-adult transition whereas most hermaphrodites of the ancestral population were still mid L4 larvae. With experimental evolution there was thus an acceleration of development from the L3-L4 moult, prior to spermatogenesis, until the L4-adult moult, when the switch to oogenesis occurs (Fig. 4B).

**Figure 4.**
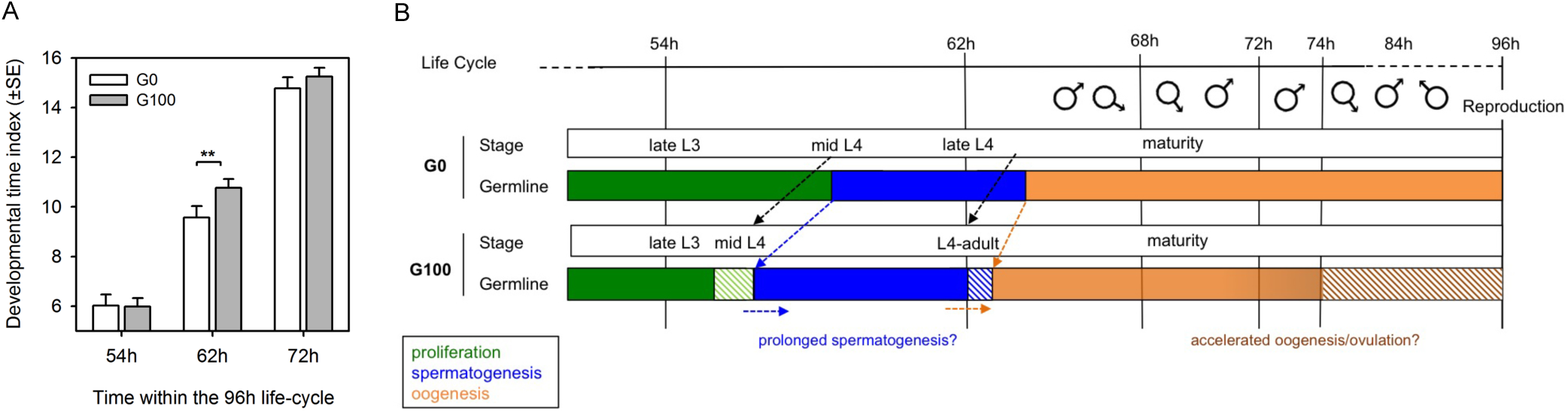
Evolution of developmental timing in soma and germline. **(A)** Quantification of developmental progression and reproductive maturation by morphological staging of a random set of individuals at 54h, 62h and 72h of the life cycle. In the ancestral population, at 54h, hermaphrodites were at the developmental stage index 6, corresponding on average to late L3 stage larvae (Table S1), and at 62h, most hermaphrodites grew to the mid L4 stage and at 72h they were all early mature adults (indices 14-15). There was no significant evolutionary response in developmental time at 54h or 72h but evolved populations were developmentally advanced at the intermediate time point of 62h. At this time point, most individuals of evolved populations had reached the L4/adult moult or early adulthood (index 11) whereas most individuals of the ancestral populations were still in the mid to late L4 stages (index 10). Linear mixed effects and generalized linear mixed effects models used to test for differences among generation 0 (G0) and generation 100 (G100): * P-values<0.05, ** for P-values<0.01, *** for P-values<0.001. Means and error least square estimates are shown. See main text for summary statistics. **(B)** Schematic outline of evolutionary heterochronic changes in soma versus germline. Ancestral hermaphrodites start spermatogenesis at mid L4 stage, switch to oogenesis in late L4 stage, reach adulthood shortly after 62h, and initiate embryogenesis around 68h of the life cycle. Before reproduction at 96h there were therefore about 30 hours where males could mate with them (Theologidis et al. 2014). Dashed arrows and boxes indicate the measured developmental changes after 100 generations under the fixed 96h life-cycle. With experimental evolution, hermaphrodites were at the midL4 stage at an earlier time, even if they delayed entry into spermatogenesis. The switch to oogenesis was also delayed, possibly because of prolonged spermatogenesis, although they reached the L4 to adult moult earlier than ancestral hermaphrodites and eventually reached maturity (onset of embryogenesis) at the same time (68h to 72h of the cycle). Oogenesis and ovulation rates were accelerated until reproduction at 96h, possibly due to the presence of males, but it is possible that this acceleration started soon after the sperm-oocyte switch.

At 72h, the great majority of hermaphrodites in both evolved and ancestral populations were young adults that had not yet started laying embryos, and so we distinguished differences in reproductive maturity by scoring the absence/presence of oocytes and counting the number of embryos *in utero*. At this time point, evolved and ancestral hermaphrodites were on average at the same developmental stage, with most individuals having one to five embryos present in the uterus, which suggests that timing of reproductive maturity, defined as onset of embryo production, did not evolve (Fig. 4A; t_16.3_ P=0.25). This results further confirm the results on an earlier measure of reproductive maturity scored at 68h (Fig. 3B). Evolution of accelerated developmental time until the switch from spermatogenesis to oogenesis was thus followed by a developmental deceleration until reproductive maturation. Given that the timing of reproductive maturity did not evolve, this suggests that the evolution of increased sperm production – and resulting delay in the onset of oogenesis – was compensated for by the compression of developmental time during the last larval stage.

## 4. Discussion

In this study we have taken advantage of *C. elegans* experimental evolution to characterize the developmental changes, in particular heterochrony, that underlie the evolution of improved hermaphrodite self-fertility and associated changes in hermaphrodite sex allocation. In our experiments, heterochrony was thus defined by differences in relative timing of developmental events in hermaphrodites of the evolved populations when concurrently compared to hermaphrodites of the ancestral population in the context of the defined 96h experimental life cycle.

Our analysis uncovers multiple heterochronic events during the evolution of hermaphrodite development, not only in gametogenesis, but also in the relative developmental progression of germline versus soma (Fig. 4B). Specifically, we found that ancestral hermaphrodites started spermatogenesis at the mid-L4 stage, switched to oogenesis in the late L4 stage and reached reproductive maturity between 68h-72h of the cycle. In contrast, evolved hermaphrodites were at the mid-L4 stage at an earlier time during the life-cycle, even though they showed a delayed entry into spermatogenesis. The switch to oogenesis was also delayed in evolved hermaphrodites, likely because of prolonged spermatogenesis; nevertheless, evolved hermaphrodites reached the L4-adult moult at an earlier time point of the life cycle than ancestral hermaphrodites. Eventually, ancestral and evolved hermaphrodites reached reproductive maturity at the same time. Subsequently, observed increases in oogenesis/ovulation (measured at 74h and 86h) of evolved hermaphrodites can explain their increased early-life fertility relative to ancestral hermaphrodites. Overall, there was little evidence for the evolution of size-related traits (germline cell precursor total numbers, gamete and embryo size), that could suggest the possible evolution of more efficient resource acquisition to both spermatogenesis and oogenesis.

Our results differ from another experimental evolution study using *C. elegans* male-hermaphrodite populations showing that an adaptive shift towards earlier self-reproduction was partly due to accelerated gametogenesis and earlier reproductive maturity (Anderson et al. 2011). It is unclear whether in Anderson et al. (2011) hermaphrodite sex allocation evolved, as evolutionary changes in hermaphrodite cross-fertility or sperm production were not measured. One explanation for the differences in developmental and reproductive timing between studies may be reduced selection under selfing (Anderson et al. 2011) since male frequencies were maintained at around 40% (Anderson et al. 2010), being thus closer to effective dioecy than in our evolution experiments where males were stably maintained at around 25% (Teotonio et al. 2012). If this was the case, then selection under outcrossing could have resulted in a general acceleration of hermaphrodite gametogenesis and developmental time to maturity, which would have improved hermaphrodite reproduction under both selfing and outcrossing. Interestingly, however, our results show that soma-germline heterochrony (Fig. 4B) allowed for the evolution of increased sperm production without delaying reproductive maturity – contrary to what is expected from observations in *C. elegans* mutants with increased or decreased self-sperm production (Hodgkin and Barnes 1991; Murray and Cutter 2011). This result shows that an apparent fitness trade-off due to the developmental coupling between spermatogenesis and oogenesis can be minimized through compensatory changes in developmental timing of both germline and soma.

*C. elegans* shows natural genetic variation in diverse hermaphrodite traits associated with reproductive and sexual functions (Hodgkin and Doniach 1997; Petrella 2014; Poullet et al. 2015), including genetic variation in temporal expression dynamics of spermatogenesis‐ and oogenesis-specific genes (Francesconi and Lehner 2014). However, it remains unknown how such variation at the gene expression level affects particular developmental traits, such as the timing of the sperm-oocyte switch, and which specific loci may be targeted by selection to generate heterochrony. Transcriptomic analyses of the complex temporal dynamics of gene expression profiles associated with germline versus somatic development provide many potential candidates (Thoemke et al. 2005; Spencer et al. 2011). While it is possible that such loci directly regulate the central germline sex determination pathway, e.g. the *fog-2* locus (Schedl and Kimble 1988), additional loci may also act indirectly on germ cell proliferation and differentiation, for example, through a variety of possible soma-germline interactions, in particular, interactions mediated by the somatic gonad (Ellis 2010; Korta and Hubbard 2010). Whether regulatory changes in genes of the so-called “heterochronic pathway”, whose mutational disruption causes pronounced temporal shifts in the progression of certain somatic *C. elegans* cell lineages (Resnick et al. 2010), could preferentially contribute to the evolution by heterochrony remains to be tested, not only in our experimental paradigm but in general.

How does heterochrony underlie the adaptive maintenance of partial selfing in our evolution experiments? Teotonio et al. (2012) has shown that evolved populations exhibited improved male competitive performance, possibly due to sexual selection; on the other hand, hermaphrodite fertility was improved specifically under selfing (Carvalho et al. 2014b), thus in part explaining the maintenance of partial selfing. From the time of reaching adulthood until reproduction in our life-cycle (from approximately 63-66h until 96h), males had ample opportunity to find and mate with hermaphrodites (Fig. 4A) (Theologidis et al. 2014). Since after insemination, larger male sperm must crawl through the hermaphrodite uterus to reach the spermatheca, where it actively displaces the smaller self-sperm to ensure cross-fertilization (Ward and Carrel 1979), our results suggest that the increased embryo retention *in utero* and the increased numbers of self-sperm, without changes in time to reproductive maturity, may have strained cross-fertilization by males. Consistent with this hypothesis, populations with higher self-sperm numbers in the spermatheca and with higher embryo retention were also those where male sperm tended to be larger (Fig. S2). The evolution of developmental timing shifts, without generating differences in the time of reproductive maturity, may have been selected because earlier reproductive maturity would provide more time for males to mate with hermaphrodites, thus limiting self-reproduction.

Pollen discounting in angiosperms, an analogous situation to the lack of outcrossing among *C. elegans* hermaphrodites, has been proposed to explain the maintenance of partial selfing (e.g. Holsinger 1991; Porcher and Lande 2005). But there is little evidence that heterochrony can underlie partial selfing through selection for optimal sex allocation although this has been – more or less explicitly – proposed for a long time (Darwin 1877; Mazer et al. 2004; Goodwillie et al. 2005; Weeks et al. 2006; Johnston et al. 2009; Escobar et al. 2011). Besides the studies in *C. elegans* mentioned (Anderson et al. 2011; Murray and Cutter 2011), a study in the monkeyflower *Mimulus guttatus* has shown that five generations of selection under restricted cross-pollination resulted in the evolution of increased corolla sizes but reduced anther-stigma distances, implying heterochrony in organ proliferation and differentiation (Roels and Kelly 2011). Similar responses were nevertheless obtained when cross-pollination was not restricted, though not to the same extent, suggesting that heterochrony caused the evolution of more efficient acquisition of resources towards reproduction instead of causing the evolution of sex allocation. In a recent study in the snail *Physa acuta*, 17 generations under restricted outcrossing resulted in reduced developmental time to maturity, although allocation towards male and female functions showed weak responses (Noel et al. 2016). Thus, the few experimental studies suggesting a potential role of heterochrony in the maintenance of partial selfing could not clearly show that heterochrony was due to the evolution of sex allocation.

In conclusion, our study shows that heterochrony in germline and soma underlies the evolution of higher reproductive success under selfing in order to counterbalance the evolution of increased reproductive success under outcrossing – in our case by the improvement male performance (Teotonio et al. 2012). Heterochrony therefore represents a specific mechanism that may be involved in many instances of partial selfing in natural populations with mixed reproduction modes (Goodwillie et al. 2005; Jarne and Auld 2006; Weeks et al. 2006). From a broader perspective, our study shows how subtle, yet complex heterochrony can underlie the adaptive evolution of major organismal traits, such as sex allocation and life history. Integrating quantitative developmental analysis with experimental evolution can thus help to understand how natural selection operates on the interplay of developmental parameters that determine fitness at the population level, a central goal of evolutionary developmental biology.

## Data Accessibility

Supplementary material accompanies the manuscript. All raw data is provided in Table S2.

## Author Contributions

N.P., A.V., C.G. and C.B. did the assays and preliminary analysis. S.C. cultured the populations for developmental characterization. H.T. and C.B. designed the project, analysed the data and wrote the manuscript.

## Acknowledgements

We thank Céline Ferrari and Nicolas Callemeyn for technical assistance, Judith Kimble for providing the RME-2 antibody, Asher Cutter for comments on a previous version of the manuscript, and M.A. Félix. P. Jarne and P. C. Phillips for discussion.

